# Neural Mechanisms of Mutualistic Fish Cleaning Behaviour: a Study in the Wild

**DOI:** 10.1101/2024.06.12.598765

**Authors:** Daniele Romeo, Sandra Ramirez-Calero, Timothy Ravasi, Riccardo Rodolfo-Metalpa, Celia Schunter

## Abstract

One crucial interaction for the health of fish communities in coral reefs is performed by the cleaner fish by removing ectoparasites and other particles from the body of other fish, so called clients. Studying the underlying mechanisms of this behaviour is essential to understanding how species react to social stimuli and defining the drivers of mutualism. Here, we pinpoint the neural molecular mechanisms in the cleaning behaviour of *Labroides dimidiatus* in the wild through an *in-situ* interaction experiment at a coral reef in New Caledonia. Five cleaners and clients (*Abudefduf saxatilis*) were placed into underwater aquaria to interact, while five were not presented with a client. The brain transcriptomes revealed 291 differentially expressed genes in cleaners that were interacting with a client. Among these genes, *grin2d, npy, slc6a3* and immediate early genes (*fosb*; *fosl1*; *nr4a1)* were related to learning and memory, glutamate and dopamine pathways, which confirm molecular pathways observed in laboratory studies. However, a new potential mechanism was found with *npy* (Neuropeptide Y) as a driver of feeding behaviour. These results show that in-situ experiments are essential for corroborating interpretations inferred from experiments in captivity and identify drivers of interspecific interactions.

## Introduction

Social behaviour affects individual fitness in animal populations, influencing their persistence (1). The interaction between two species can lead to conflictual situations, such as competing for the same limited resources (competition) but can also be a beneficial collaboration (mutualism) where both parties gain an advantage from the interaction (2). Over time, the expression of such interaction behaviours has been shaped by natural selection by finetuning the ability of individuals to maximize the benefits of social interactions (3). This led some species to specialize and evolve sophisticated social behaviours, using interaction with other individuals to cover basic biological needs such as feeding or predator defence (4,5). Mutualistic interactions are frequent in nature and require a high grade of social skills and cognition of the social environment (5–7). Thus, species involved in a mutualistic interaction can exploit the most marginal environments, capitalizing on unoccupied niches and avoiding competition (8). Therefore, mutualisms are drivers of ecosystem complexity and functions (9).

In coral reefs, the blue streak cleaner wrasse *Labroides dimidiatus* relies on social interactions as its main trophic source (10). Its survival is based on the ability to clean other fish (called clients) consuming parasites, mucus, and dead skins from their bodies (11). This cleaner fish can have over 2200 interactions per day and is considered a “dedicated cleaner” due to its persistence in cleaning throughout its life (12). Due to the central role of it social interactions, the cleaner wrasse has evolved decision-making, reputation management, and social skills to better manage its reputation and clientele in every social context (13,14). Indeed, *L. dimidiatus* has the capacity to prioritize different clients based on their ecological patterns (accessibility to only one or multiple cleaning station within the home range) and adjust cooperation levels (the ratio between parasites and mucus eaten) in the presence of a bystander client fish (4,14). Therefore, the ability to use social skills and the phenotypic flexibility shown in cleaning interactions make *L. dimidiatus* an ideal species for examining the drivers of such social interaction behaviour.

Gene expression is one of the main processes involved in the phenotypic response to the social environment, modulating species’ plasticity in the short- and long-term (15,16). For instance, differential expression of Vitellogenin drives the division of labour in Hymenoptera (16) and knocking out the Hrh1 gene reduces aggressiveness in mice and zebrafish when exposed to an unfamiliar individual (17,18). For cichlids, *Astatotilapia burtoni*, behavioural responses to intruders or gravid females are regulated by nonapeptides and sex steroid gene expression (19). For the cleaning behaviour in *L. dimidiatus*, glutamatergic receptors, immediate early genes (IEGs), isotocin, estrogen and progesterone receptors as well as dopaminergic pathway genes play a central role in the brain (20). Processes of learning and memory in the cleaner wrasse are suggested to be mediated by glutamate through different expression of both ionotropic and metabotropic receptor genes, while partner recognition, which is a key factor influencing the interaction behaviour, may be driven by IEGs (20). In addition, reduction of dominance towards the client and promotion of prosocial behaviour have been associated with the expression of estrogen and progesterone (20). Thus, different gene expression patterns modulate the species’ response to social stimuli, influencing their cleaning success by promoting their capacity to react to the social environment.

Most of our understanding of molecular drivers underlying social behaviours comes from mechanistic experiments conducted in captivity, and the lack of in-situ experimental data is evident. Gathering data in natural settings eliminates a potential distortion in phenotypic signals caused by captivity conditions in lab-based experiments (21,22). Thus, in-situ experiments are essential to corroborate or contest conclusions observed in ex-situ experiments. Therefore, our aim is to unravel the neural molecular drivers involved in the interaction behaviour of *L. dimidiatus* and its client through an in-situ experiment in the wild. As observed in previous studies in captivity, we may expect to find differential expression of Immediate Early Genes and genes related to neurotransmitters such as dopamine, isotocin and glutamate as main molecular drivers of cleaning interaction. Understanding the underlying mechanisms of wild cleaner’s interactions with clients will elucidate the drivers that prompt two species to engage in a mutually beneficial relationship.

## Methods

### Sampling and behavioural experiment

This experiment was conducted in southwestern New Caledonia (-21.953529, 166.004627) between 12^th^ and 14^th^ March 2020. Ten individuals of *L. dimidiatus* and five individuals of the clients *Abudefduf sexfasciatus* were collected on SCUBA using barrier nets and hand-nets. The experimental set-up included three experimental tanks (20 x 20 x 30 cm) that were placed underwater on the seafloor, filled one-fifth with sand and a weight to avoid buoyancy at approximately 30 meters from the coral reef (fig. 1). A video camera, GoPro Hero 6 Black, was placed in front of each tank.

**Figure 1:**
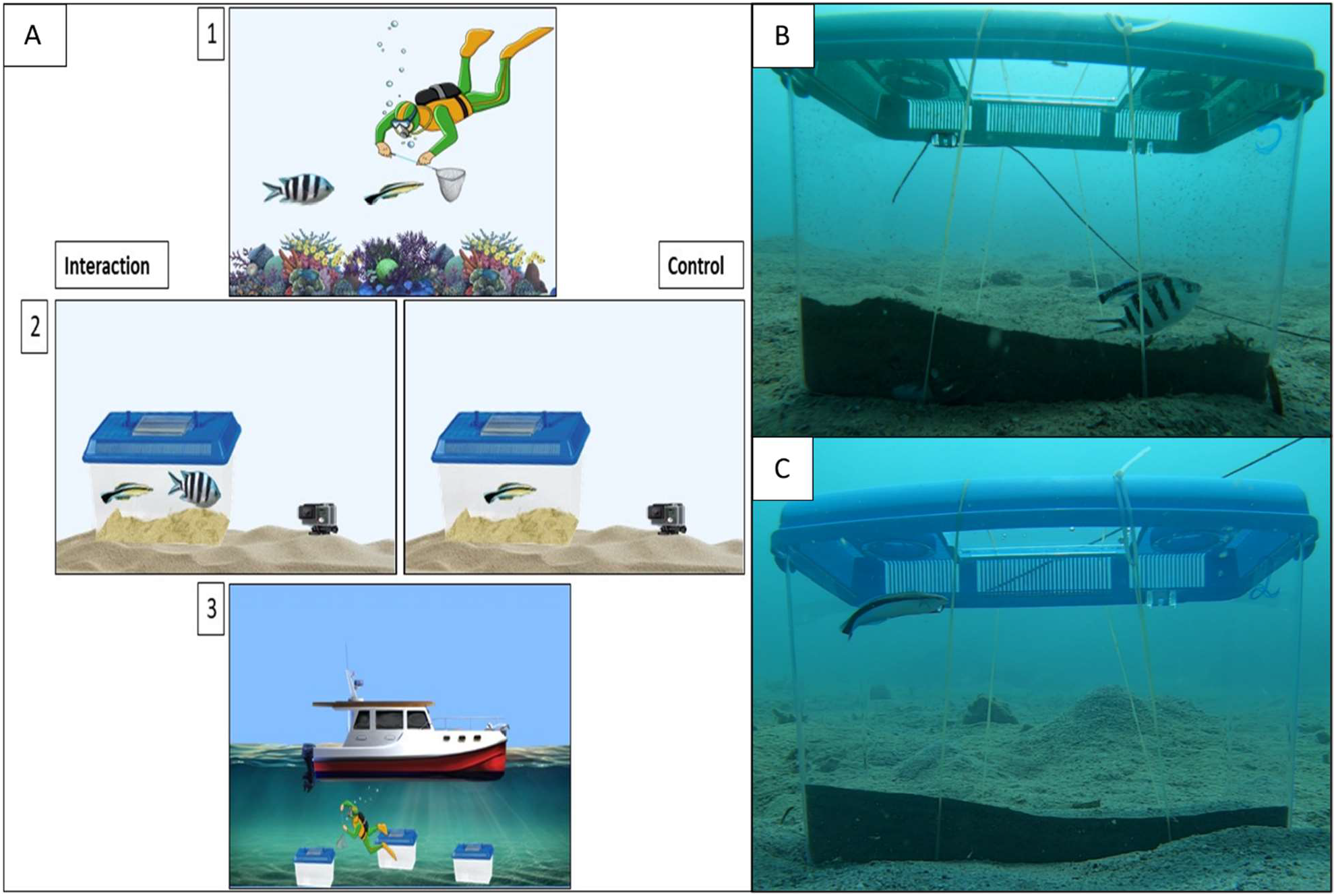
A) In-situ experiment: Labroides dimidiatus are collected and placed in aquaria underwater whether with a client (Interaction) or without (Control). After the experiment, the diver brought the cleaner to the surface for brain collection. B) Experimental set up for both condition (interaction (B) and control (C)).

Collected cleaner fish were placed first into the experimental tanks, followed by the client shortly after. To isolate the gene expression signal in cleaning interaction, a control condition was adopted where cleaner fish were placed in the tanks without clients (figure 1A2). The recording was immediately started as the fish were placed in the tank, and their behaviour was recorded for 50 minutes. Directly after, the fish were collected and brought to the water surface, where the brains were dissected immediately on the boat, stored in RNAlater (Invitrogen), and frozen after 24 hours. Subsequently, the samples were shipped to The University of Hong Kong for further processing and kept in the -80° freezer.

### Behavioural analysis

To determine if the cleaner fish interacted with the client and to detect any abnormal behaviour in the control cleaner fish, videos were analysed using the software Boris v.8.22.17 (23). The first five minutes were considered as acclimatization and were not considered in the analysis. For the interaction treatment, five different behaviours, previously standardized and calibrated in laboratory conditions (20), were evaluated: Interaction, Dance, Tactile Stimulation, Inactivity and Jolt. The time of occurrence and the duration of the first four were counted in seconds; while for Jolts only the time of occurrence was counted. Analysis parameters were considered as follows: Interaction: each time the cleaner fish approaches and inspects the client or exhibits the typical behaviour of cleaning interaction such as dancing, tactile stimulation or biting the surface of the client’s body. Dance: broad and smooth symmetrical longitudinal movement to capture the attention of the client, directly linked with its willingness to interact (24). Tactile Stimulation: the cleaner places itself over the dorsal part of the client and through slightly inclination rapidly moves the pelvic fins on the back of the client (25). Inactivity: the time spent by the cleaner lying down without moving for more than two seconds, which is directly linked to stress (26). Jolt: a sudden movement of the client in response to a bite of the cleaner during an interaction which is considered cheating behaviour (breaking of the mutualism) (27).

### RNA Extraction and Transcriptome Analysis

Total RNA was extracted from cleaner fish whole brain tissue using RNeasy Mini Kit (Qiagen), and the quality and quantity was checked on a 4200 TapeStation (Agilent) and a nanodrop respectively. High-quality samples (RIN>8) were used, and cDNA libraries were prepared by KAPA mRNA HyperPrep Kit and sequenced paired-end 151bp on an Illumina NovaSeq 6000 at the Centre for PanorOmic Sciences (CPOS) of the University of Hong Kong. An average of 38 (±1.23) million reads per sample were obtained, and the quality was checked by FastQC v. 0.12.1. Illumina adapters and low-quality reads were removed using Trimmomatic v0.39 (28), using the parameters: SLIDINGWINDOW:4:30; MINLEN: 40; threads: 32; 2:30:15:8:true (surviving reads 96% (±0.13)). Subsequently, to map the reads to the reference genome (29), the software HISAT2 v.2.2.1 (30) was used adopting default parameters, with a mapping rate average of 83% (±10.05%). To count the number of reads mapped to each gene in the reference, we used FeatureCounts with default parameters (31).

To statistically assess the differential gene expression between the cleaner fish that interacted with the client and the control group (cleaner fish alone), we employed the package DESeq2 v.3.18 (32) with a Wald Test statistic while adopting FDR p-adjusted significance value of 0.05 as a cut-off. Furthermore, to analyse networks of genes with significantly correlated expression patterns, weighted correlation network analysis was carried out with WGCNA v1.72-5 (33). Gene module networks were created with the command blockwiseModules by using “signed” as topological overlap measure (TOM) by setting 2000 and 30 as the cut-off for the highest and lowest number of genes in the network and an elevated power value of 21 due to the relatively low sample size. Then, eigengene values for each module were correlated with the time (in seconds) spent by the cleaner interacting with the client by using the Pearson correlation. The expression patterns of the modules of which eigengenes resulted significantly correlated with the trait (p-value < 0.05) were further analysed with paired t-tests between interaction and control. Functional enrichment on the DEGs and on the genes of the modules with significantly correlated expression patterns between interaction and control was carried out by using OmicsBox v. 3.1 using Fisher’s exact test with a cut-off of FDR 0.05 (34).

## Results

### Behavioural analysis

The cleaner fish placed in the tank with the client on average spent 32.5 ± 20% (1069 ± 672 seconds) of the time interacting and 8 ± 18% in an inactivity state. Behavioural results are reported in detail in Supplementary Table 1. Control cleaner fish did not show any abnormal behaviour, except for one fish, which showed stereotyped movement (recurring circular displacements) and long inactivity time (2048 seconds; 39% of the total time). Therefore, this fish LD9 was excluded from the transcriptomic analysis.

### Transcriptomic analysis

The cleaner fish interacting with the client exhibited 291 differentially expressed genes (DEGs) compared to the cleaner fish in the control group (Supplementary Table 2). In total, 154 enriched functions emerged from these DEGs, 114 of which were biological processes (Supplementary Table 3). Several differentially expressed genes (*gria4, grin3a, grin2d*) were related to glutamatergic synapse, and glutamate-gated receptor activity GO terms (fig. 2A, Supplementary Table 3). Several immediate early genes (IEGs) (*fosl1, fosb, foxo1a, nr4a*(*1 & 3*), *egr2b*) (fig. 3) and second cellular messengers such as *zfp36l3, cnga3* and *map4k2* related to learning and memory processes were also differentially expressed (fig. 2B, Supplementary Table 3). Dopamine was further altered by the interaction behaviour, reflected in dopamine biosynthetic process and regulation of dopamine metabolic process with genes *slc6a3, npy, nr4a1* and *nr4a3* underlying these functions (fig. 2C). Furthermore, the gene *npy* encoding for neuropeptide Y is involved in feeding behaviour function among other genes (fig. 2D; fig. 3). Considering the WGCNA analysis, 193 gene modules were produced, of which 12 significantly correlated with the interaction trait (Supplementary Table 4, Supplementary Figure 1). Functional enrichment analysis produced results for two modules (Sky Blue and Dark Red) (Supplementary Figure 2, 3). The module Sky Blue showed enrichment in functions related to tRNA processing and regulation of chromosome segregation (Supplementary Table 5; Supplementary Figure 2), while the module Dark Red was related to metabolism, such as lipids, protein, pyrimidine-containing compounds, regulation of glucose import and catabolic processes (Supplementary Table 6; Supplementary Figure 3). Metabolism was also a predominant function of genes in two other modules such as Powder Blue (*apoo, ddhd1, trmt61b* and *pgs1*) and Chocolate (*bdh1, mmut*). Genes in the Powder Blue module are also related to glutamate pathway (*frrs1l, dglucy*), neurotransmitter GABA (*slc6a13*), glucocorticoids (*gmeb1*), vision (*opn3*) and epigenetic processes (*setd7*) while in the module Chocolate genes are involved in synaptic plasticity *(arf1 and sstr5)* and circadian rhythm *(prkg1, per2, and nfil3)* (Supplementary Table 4).

**Figure 2:**
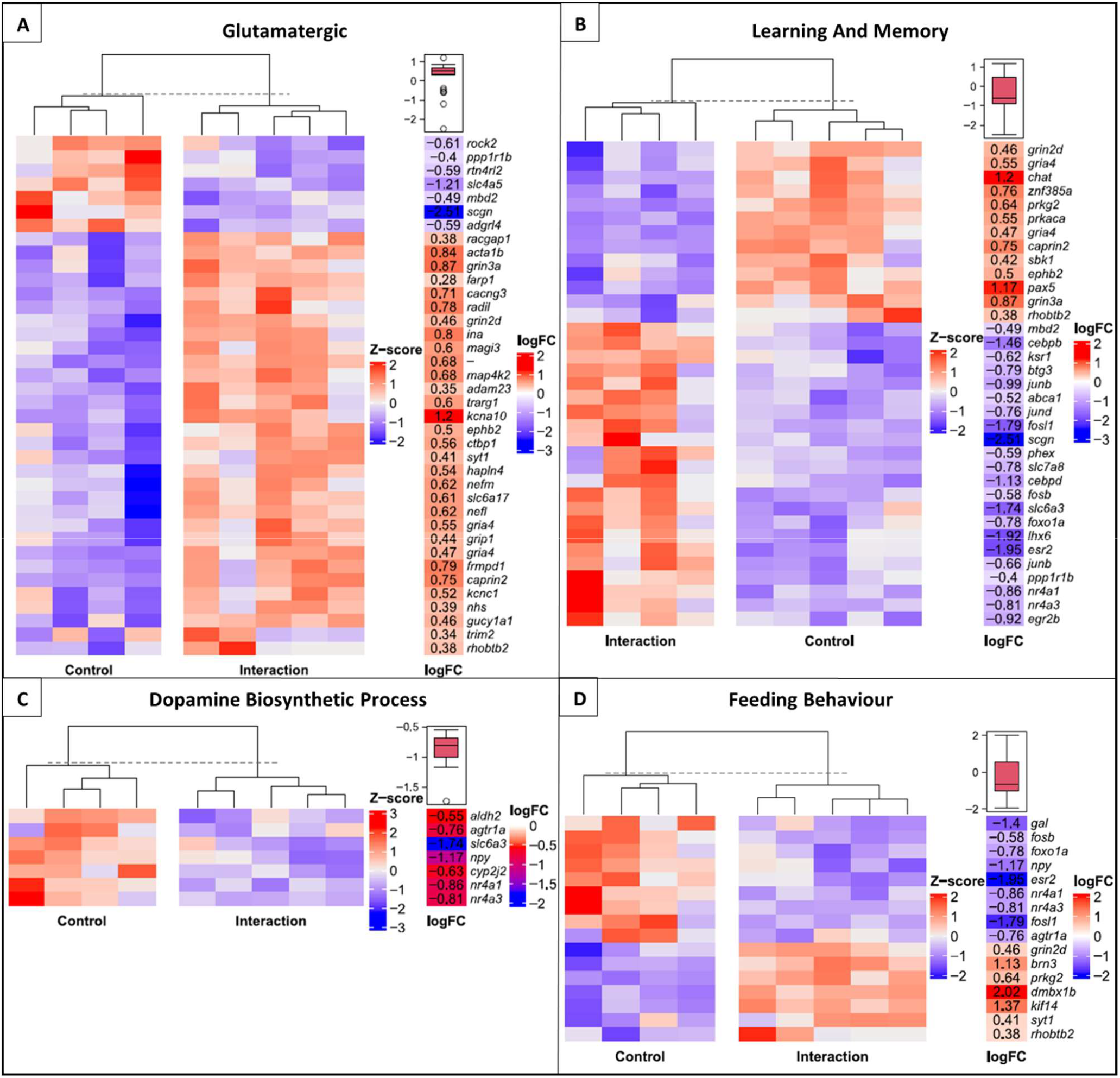
Enriched functions in the differentially expressed genes between cleaner fish that were not exposed to the client (control) and cleaner fish that were exposed (Interaction) for A) glutamatergic synapse, B) learning and memory function, C) dopamine biosynthetic processes and D) feeding behaviour. For each enriched function genes names are indicated and the Z-score related to the heatmap indicates the level of differential expression of each gene, while the column logFC refers to the log2fold change of each gene. The boxplot above logFC column shows the average of the gene expression related to the function in the cleaner fish that interacted with the client.

**Figure 3:**
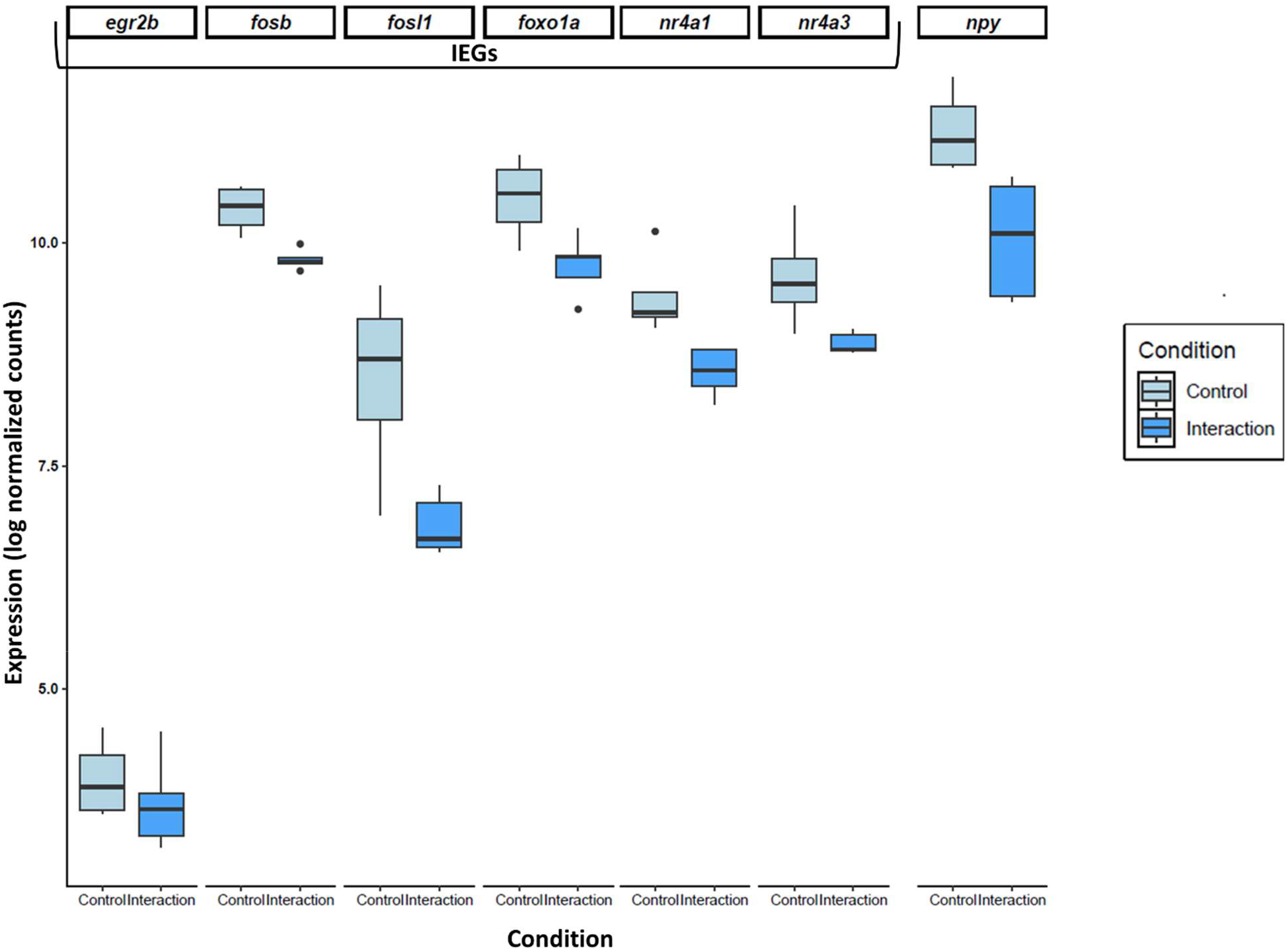
Differential gene expression of Immediate Early Genes (IEGs) and npy gene. In light blue the control cleaner fish and in blue the cleaner fish that interacted with the client. The differential gene expression is showed as log normalized counts.

## Discussions

This study reveals the transcriptional drivers in *L. dimidiatus* when interacting with a client in the wild. The interaction behaviour provoked changes in genes related to glutamate pathway, immediate early genes and dopamine pathways, but did not detect changes in genes related to isotocin as found in studies in captivity (20). However, we discovered novel gene expression changes, such as the neuropeptide Y. Therefore, our study corroborates previous findings in *L. dimidiatus*, but also adds additional molecular drivers underlying the interaction behaviour.

We found differential expression related to glutamatergic synapsis and glutamate receptor activity that mediate learning and memory processes, confirming previous findings on this species in which glutamate is one of the main molecular drivers in the cleaning interaction (20). Glutamate ionotropic receptors (NMDA and AMPA) are known to drive the social recognition memory consolidation after a social stimulus in rats (35), improving their social recognition potency of familiar or unfamiliar individuals when treated with NMDA (36). For *L. dimidiatus*, the upregulation of genes related to these receptors (*gria4, grin3a, grin2d*) may allow the recognition of clients, which is a key feature as it interacts over 2000 times daily and can adjust the outcome of the interaction through partner prioritization or adjusting service (13,37). Moreover, we found upregulation of gene *grip1* in the cleaners that interacted with the clients, which encodes for AMPAR-binding protein GRIP1 and promotes synaptic plasticity by inserting AMPARS into synapses that may enhance learning and memory processes (38), also important for cleaning interactions. Furthermore, synaptic plasticity and circadian rhythm genes showed correlated expression patterns with clock genes being able to influence neuronal activity and excitability, affecting memory consolidation and recalling learned behaviour (39,40). Therefore, no matter if in a wild or a laboratory setting, glutamatergic pathways are one of the main molecular drivers in the cleaner wrasse brain, and together with clock genes, regulate synaptic plasticity and learning and memory processes.

Immediate Early Genes (IEGs) are modulators of social behaviour in the social brain network, a highly conserved neural network in the telencephalon and diencephalon (41), and were differentially expressed when the cleaner interacted with the client. These genes are the first that react to extracellular stimuli and are associated with neuronal activity (*fosl1, fosb, egr2b, foxo1a*) and neural plasticity (*nr4a1, nr4a3*) that affect social behaviour (42–45). They drive the social decision-making in *A. burtoni*, whether to cooperate with another male to defend his territory from an intruder or to exploit the social opportunity to ascend as a dominant male (19,46). Furthermore, the processes involving IEGs are mediated by second cellular messengers such as mitogen-activated protein kinase (MAPK) and cAMP pathways, as shown in the fighting fish *Betta splendens*, where the brain-transcriptomic changes during a fight with a conspecific are associated with IEGs and MAPK pathway genes (47). *Labroides dimidiatus* upregulated the gene *map4k2*, amongst other genes involved in cAMP pathways such as *zfp36l3* and *cnga3*. Therefore, when exposed to social stimuli, IEGs together with cAMP and MAPK pathways could synergistically lead to a downstream molecular response cascade that can drive the decision-making process of *L. dimidiatus* on whether to exploit the social opportunity to approach the client and choose which behaviour to perform (dancing, cleaning, tactile stimulation or cheating). Thus, IEGs mediate *L. dimidiatus* cleaning behaviour in the wild, influencing its decision-making on whether and how to interact with the client.

Genes involved in dopamine pathways were also altered with the interaction behaviour in *L. dimidiatus*. An important regulator of synaptic dopamine availability is *DAT1* (Dopamine Transporter 1) encoded by *slc6a*, which was downregulated in interacting cleaner wrasses. Lower expression of *slc6a*, which is involved in the uptake of dopamine and extracellular clearance, would suggest higher dopamine levels in cleaner fish that interacted with the client (48). Induction of higher dopamine levels in rats or primates increases social interactions and an exaggeration of behaviours related to social rank, such as subordinates becoming more subservient in their social interactions (49,50). During the cleaning interaction, the client may impose such partner control mechanisms (chasing the cleaner) to avoid cheating behaviour by the cleaner, and therefore, downregulation of *slc6a3* may show submissive behaviour in the cleaner fish through increasing extracellular dopamine levels. Furthermore, pharmacologically blocking the D1 and D2 receptors promotes the willingness of the cleaner to interact with the client and provide tactile stimulation (51). Here, we detected changes in genes (*nr4a1* and *nr4a3*) that are linked to D1 and D2 receptor activity. In fact, in mice, overexpression of *nr4a1* impairs D1 and D2 receptor signalling (the effect of *nr4a3* is still unclear) (44,45). Therefore, changes in *nr4a(1*,*3)* could alter D1 and D2 receptor pathways and regulate cleaner fish behaviour by mediating submissive behaviour via dopamine extracellular concentrations. Thus, the dopaminergic pathway is another molecular driver of cleaner fish interspecific social behaviour in the wild.

Interestingly, *npy*, a gene encoding for neuropeptide Y (NPY) known for its role in food intake (52–54), was downregulated in the fish that interacted with the client. NPY is one of the brain’s most abundant and effective orexigenic peptides (53). In sturgeon fish, for instance, *npy* brain expression decreased after a meal and in goldfish, injection of Y1 and Y5 receptor agonists increased food intake while food deprivation increased hypothalamic expression of *npy* mRNA(55–57). Hence, fasting leads to an increase in *npy* expression promoting food intake behaviour, while following a meal, expression levels decrease in numerous teleost fish (58–61). In fish, NPY can negatively affect food intake behaviour by inhibiting dopamine neurons through pre-synaptic and post-synaptic mechanisms in the ventral tegmental area (VTA), the brain region where the mesocorticolimbic dopamine system controls food intake, food reward and feeding-related behaviours (62). Therefore, we may hypothesize that upregulated expressed of *npy* in the control cleaner fish, could drive to seek out clients and initiate cleaning interactions, while a low level of *npy* can prevent cleaner fish from interacting, modulating the mesocorticolimbic dopamine system through food reward processes. Moreover, the differences in genes related to metabolism observed in gene networks correlated with the interaction behaviour show support for physiological mechanisms triggered after a meal. Thus, overexpression of *npy* may be a promoter of cleaning interaction in *L. dimidiatus*, making this gene one of the molecular drivers in cleaning behaviour.

The brain transcriptomic profile involved in cleaning interaction found in this study corroborate previous studies in *L. dimidiatus* cleaning behaviour where glutamate, IEGs, and dopamine were shown to be molecular drivers. Moreover, we indicate *npy* as an additional regulator of this interspecific interaction revealing that while in-situ studies provide a mechanistic approach to studying interactions, ex-situ experiments allow important additional insights into wild animal behaviour. The molecular processes in the cleaning interaction direct the regulation of partner recognition and memory consolidation to recognize clients and retain information for future interactions. This allows social decision-making on whether and how to interact with the client, feeding and submissive behaviour to promote interspecific interaction and honesty while cleaning. All these features, driven by the molecular pathways, affect the efficiency of cleaner fish behaviour and thus the persistence of one of the most crucial interspecific interactions for the balance and health of the coral reef ecosystem.

## Data accessibility

The raw sequencing data can be found NCBI Bioproject number PRJNA1120171. Reviewer link can be accessed here: https://dataview.ncbi.nlm.nih.gov/object/PRJNA1120171?reviewer=n10bqpe73p6t48tsvn5r4qvi23

## Authors’ contributions

C.S. designed the experiment and sample collection was conducted by S.R.C., C.S., T.R. and R.R-M. D.R. performed RNA extractions, analyzed the behavioural videos and transcriptomic data with input from C.S. D.R. and C.S. wrote the first draft of the manuscript and all the authors revised and approved the final manuscript.

## Acknowledgments

We are thankful to Erina Kawai, Michael Izumiyama and Billy Moore for aiding with the sample collection. We want to acknowledge the Schunter lab members at The University of Hong Kong Dr. Sneha Suresh, Arthur Yan Chi Chung, Jade Sourisse, Maxine Cutracci, Dr. Lucrezia Celeste Bonzi and Maddalena Ranucci for their feedback and comments provided to this work. We also thank our collaborator Dr. José Ricardo Paula, for the stimulating discussions on the topic and the feedback on data analysis.

## Ethics statement

The experiment was performed under the permits granted from Province Sud (New Caledonia), project SuperNatural N. 34314-2019/3-REP/DENV.

