## SupplementaryFigures for "Neural Mechanisms of Mutualistic Fish Cleaning Behaviour: a Study in the Wild"

### Supplementary Figures

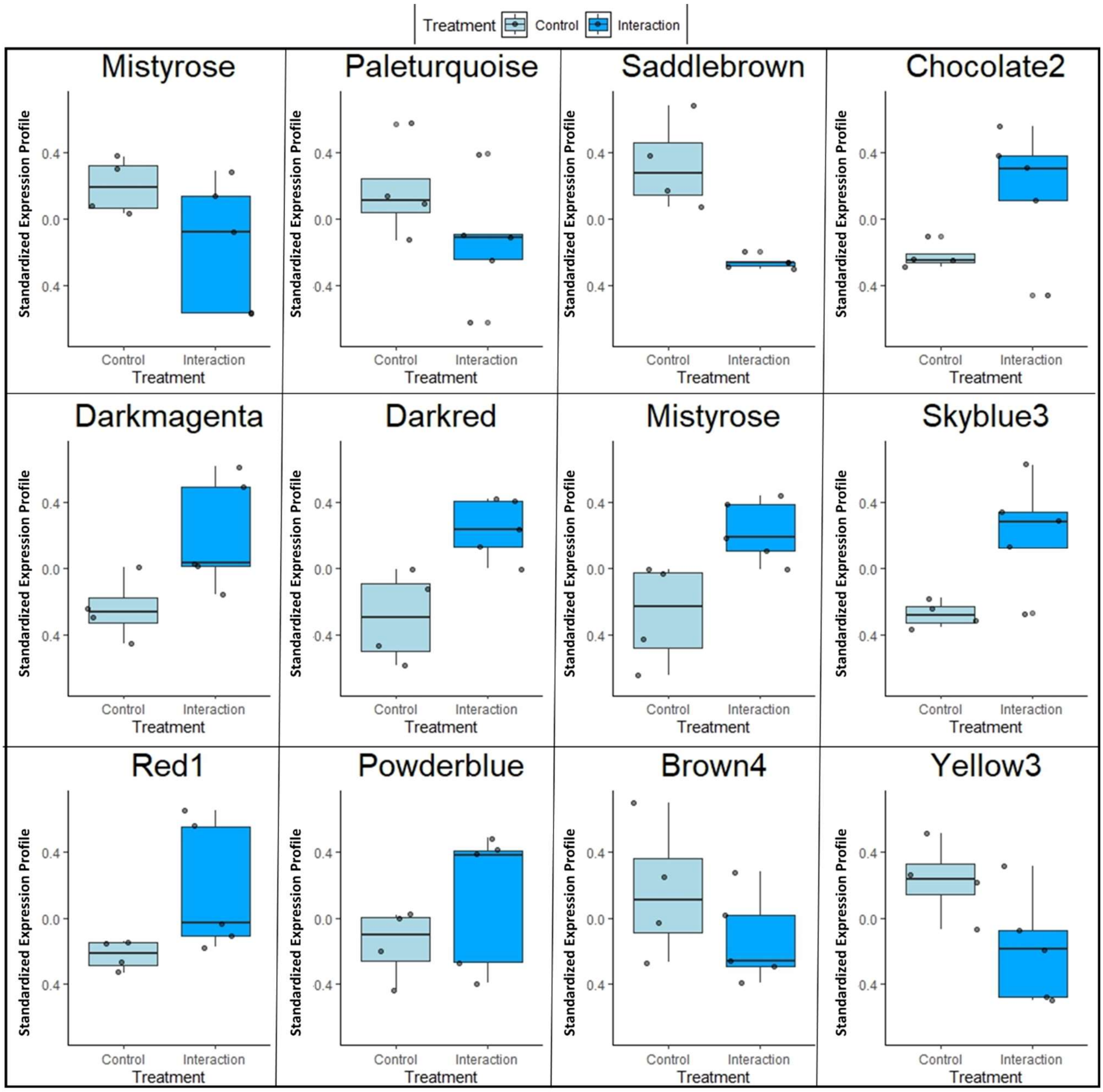


***Supplementary Figure 1****: Boxplots showing the standardized expression profile after the detection of the module eigengene in WGCNA analysis, in control cleaner fish (light blue) and cleaner fish that interacted with clients (blue).*

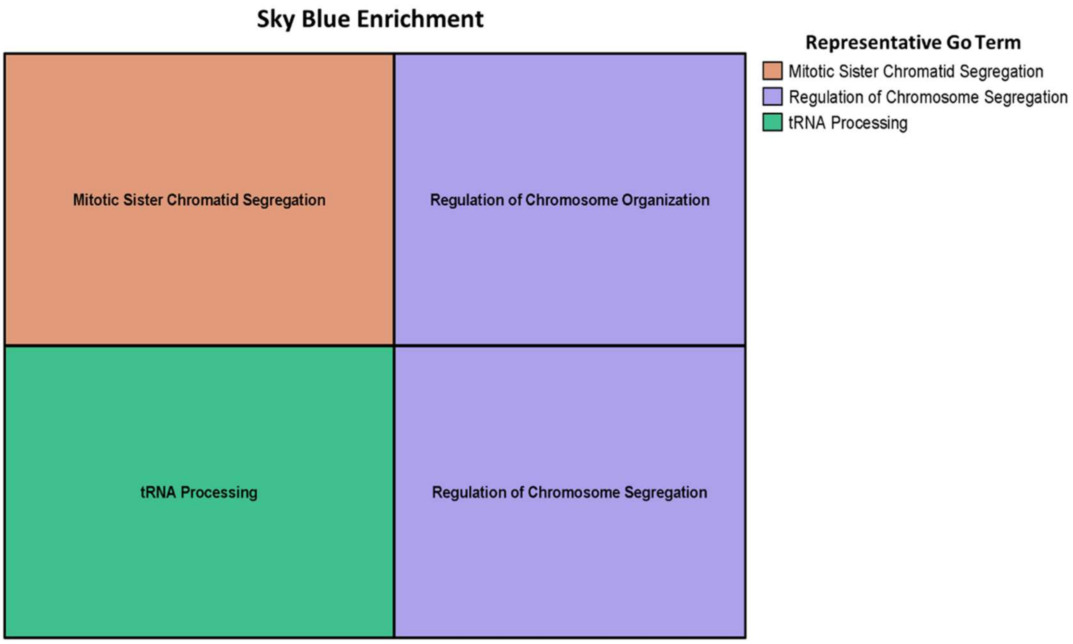


***Supplementary Figure 2****: Enriched functions related to the module "Sky Blue". In each box there are the descriptions for each go term, while the colours indicate the representative go terms.*

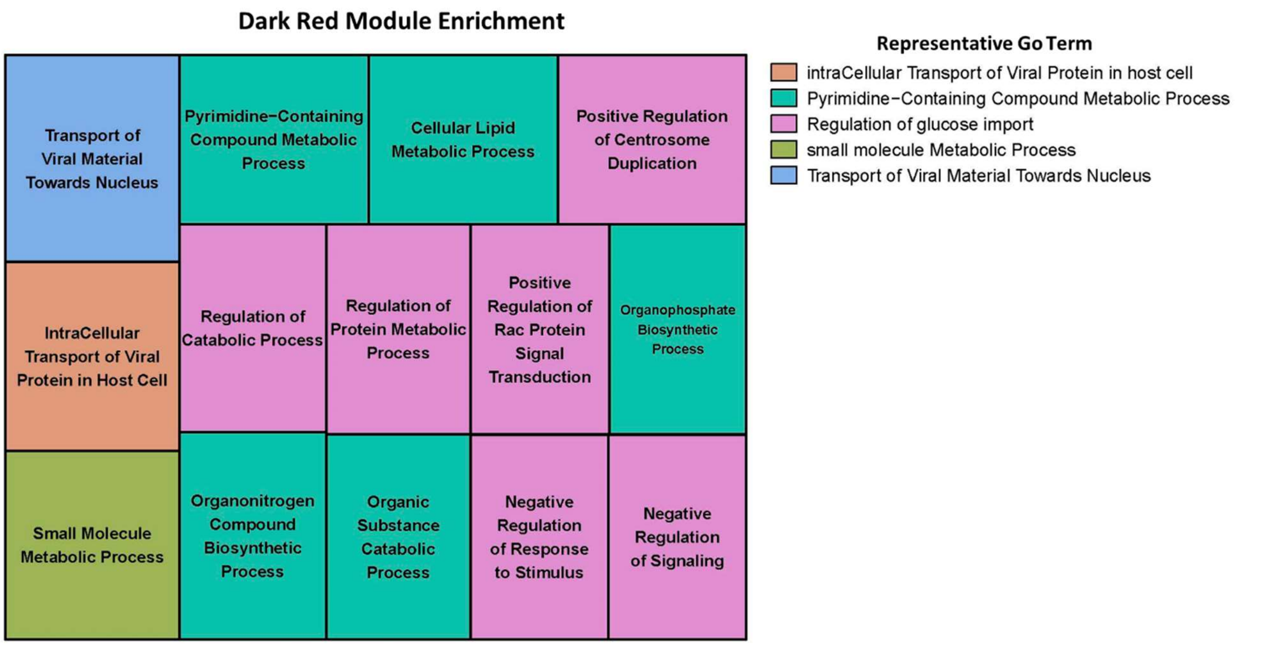


***Supplementary Figure 3****: Enriched functions related to the module "Dark Red". In each box there are the descriptions for each go term, while the colours indicate the representative go terms.*
